# High-resolution lac insect genome assembly provides genetic insights into lac synthesis and evolution of scale insects

**DOI:** 10.1101/2023.02.05.526168

**Authors:** Weiwei Wang, Xiaoming Chen, Nawaz Haider Bashir, Qin Lu, Jinwen Zhang, Xiaofei Ling, Weifeng Ding, Hang Chen

**Affiliations:** Institute of Highland Forest Science, Chinese Academy of Forestry, Kunming 650224, China; The Key Laboratory of Cultivating and Utilization of Resources Insects, State Forestry Administration, Kunming 650224, China

**Keywords:** *Kerria lacca*, Isoprenyl diphosphate synthase, Lac synthesis, Horizontal gene transfer

## Abstract

Lac insect is an important resource insect with great commercial value. However, the lack of its genome information restricts the fundamental biological research and applied studies of this species. Here, we first assembled the contig of *Kerria lacca* using Illumina and Nanopore sequencing by Hi-C and T2T techniques to obtain the genome of *K. lacca* at the chromosome 0 gap level. The genome of *K. lacca* was 256.62 Mb and the Scaffold N50 was 28.53 Mb. A total of 56.94 Mb of repeat sequences, constituting 22.19% of the assembled genome, were identified. We annotated 10,696 protein-encoding geneswith 89.74% annotated. By horizontal gene transfer analysis, we obtained a putative gene Isoprenyl diphosphate synthase (IPPS), a key enzyme in the isoprene synthesis pathway in the regulation of lac biosynthesis that transferred horizontally from bacteria to the genome of *K. lacca*. Meanwhile, we constructed the lac synthesis biosynthetic pathway and screened 25 putative key genes in the synthesis pathway by transcriptome analysis in different developmental stages and tissues. It provides new research ideas to reveal the molecular mechanism of lac biosynthesis and regulation. The high-quality chromosomal-level genome of lac insect will provide a basis for the study of development, genetics, and the evolution of scale insects.

## 1. Introduction

*Kerria* spp. is a herbivorous insect of the order Hemiptera (Coccoidea, Tachardiidae), with 29 species worldwide (Varshney and Sharma, 2020; Talukder and Das 2020; Rajgopal et al., 2021; Bashir et al., 2021), and the majority of them found in tropical and subtropical regions of south Asia and southeast Asia between 70-120 °E and 8-32 °N (Varshney and Sharma, 2020). *K. lacca* has become an important insect species for producing high-quality lac because of its high lac resin content, low wax, less impurities, and light color, and it is an important forest resource insect in a country. Lac insects can parasitize over 300 host plant species, such as *Schleichera oleosa, Flemingia macrophytla* and *Cajanus cajan*, and suck host plant phloem using piercing-sucking mouthparts. Simultaneously, females secrete the secondary metabolite lac through glands (Chen and Feng, 2009).

Shellac is the only animal-secreted natural resin that has been extensively researched and utilized in a wide range of ways; its primary components are shellac resin, shellac wax, and shellac dye. Lac resin is a mixture of lactones and lactides derived from hydroxy-fatty acids and sesquiterpene esters; among them, the hydroxy-fatty acid is mainly aleutric acid, whereas the sesquiterpene acid is primarily jalaric acid (Chen et al., 2008, Chen et al., 2021). Because of its exceptional qualities such as strong adhesion, good insulation, strong plasticity, moisture resistance, corrosion resistance, and smooth coating, shellac is considered an important natural forest chemical raw resource. It is frequently utilized after processing in the military industry, chemical industry, electronics, food, medicine, and other industries with significant economic significance (Chen, 2005; Thombare et al., 2022). Moreover, due to the fact that it is natural, non-toxic, biodegradable, and good for the environment, it can also be used as an eco-friendly material and has a lot of potential for green technology. Researchers predicted that the shellac industry will require about 7,500-8,000 tons of shellac and its value-added products annually, resulting in economic benefits of 80-90 million US dollars (Thombare et al., 2022).

Extensive research on lac insects has been conducted both domestically and internationally, with a focus on their biological characteristics, ecological adaptability, host plants, genetic variety, and so on; however, research on the mechanism of lac synthesis is still lacking. In their investigation of genes associated with lac resin production, Shamim et al. discovered that acetyl-CoA is a common precursor for the synthesis of jalaric acid and sesquiterpene acid. A transcriptomic study of *K. chinensis* at different developmental stages screened 28 candidate genes related to lac synthesis, verified the function of the screened FAD gene, and found that this gene has a regulatory effect on lac secretion (Wang et al., 2019a, 2019b). At present, the rule of lac secretion by lac insect and the molecular mechanism of lac formation in the body are unclear, and effective methods to improve the quantity and quality of lac secreted by lac insect have not been discovered, which has become a barrier in the development of the lac industry both at home and abroad. Understanding the molecular regulation mechanism of lac biosynthesis is of great significance for optimizing lac insect secretion and establishing sustainable and effective lac production.

Here, we report the first high-quality genome assembly of the lac insect. In this study, we used high-quality whole genome data of the lac insect *K. lacca*, along with transcriptome analysis of different development stages and tissues, to comprehensively explore the mechanism of lac synthesis. The findings of this study have the potential to increase the genetic information of scale insects, further elucidate the molecular regulation mechanism of lac biosynthesis, and also be used as a valuable resource in the development, genetics, and evolution of Kerriidae. This lays a molecular foundation for the protection, improvement, and exploitation of lac insect genetic resources.

## 2. Materials and methods

### 2.1 Samples

Samples of Kerria *lacca* were reared on *Schleichera oleosa* at the Experimental Station of Yuanjiang, Institute of Highland Forest Science, Chinese Academy of Forestry in Yunnan Province, China (102°00′46″ E, 23°36′11″ N). Female lac insects at late adult (n = 100) stages were collected. To perform transcriptome analysis of different developmental stages and adult tissues, female lac insects larvae (n =100), early adults (n = 100), mid adults (n =100), and late adults (n = 100) were collected. Additionally, four tissue samples were obtained from females at the late adult stage (A3) using a sterilized scalpel and tweezers: anal tubercle (AT), brachia (BH), body wall (BW) and midgut (MG). Reactions were carried out in three replicates for each biological sample. Immediately, all samples were put in liquid nitrogen, then transferred and stored at -80 °C. The samples were used for DNA and RNA extraction, library construction, and sequencing.

### 2.2 Genomic sequencing

Genomic DNA (gDNA) was isolated from 100 individual female lac insects at late adult stages using the CTAB method, following the manufacturer’s instructions. After measuring quality and quantity, a library with an average insert size ∼150 bp was constructed and sequenced using the Illumina NovaSeq 6000 (Illumina, San Diego, CA, USA) platform. For third-generation sequencing, a library with an average insert size ∼15 Kb was constructed. The libraries were sequenced on the ONT (Oxford Nanopore Technologies, Oxford, UK) Sequel platform with Sequel PromethION P48. For Hi-C analysis, 100 lac insect specimens were soaked in 1% formaldehyde for 10 min at room temperature and in a 2.5 M-glycine solution to terminate the isolation and cross-linking of lac insect cells. After adaptor ligation and quality control, the Hi-C library was sequenced using the Illumina HiSeq PE150 platform.

### 2.3 Genomic assembly

The Illumina paired-end reads were used for 19 bases k-mer analysis to estimate the genome size and heterozygosity. To construct the chromosome-level genome assembly, the Hi-C sequencing data were aligned to the final assembled contigs by ALLHiC (v0.9.12) and juicer (v1.6) (Neva et al., 2016b) to obtain the interaction matrix, and then the contigs were sorted and anchored using 3D-DNA (v180419) (Dudchenko et al., 2017). Finally, the Hi-C contact maps were applied to the 3D-DNA visualization module and reviewed with juicebox Assembly Tools (v1.11.08) (Neva et al., 2016a) for quality control and interactive correction of the automatic output. A chromosome-level genome sequence was obtained by using 100 N complement gaps from sequenced, directed, and non-redundant contig sequences. For genome assembly, the clean Nanopore reads after filtering and decontamination were assembled with Next Denovo (https://github.com/Nextomics/NextDenovo/releases/tag/v2.3.1). To fix sequencing errors in the final assembly, Racon (version: 1.4.11) and Pilon (version: 1.23) were chosen to polish the genome using long sequencing data and short reads with default parameters for two rounds each. In order to obtain a 0 gap genome, we respectively combined the ONT raw data and wtdbg assembly method to obtain a contig level genome using winnowmap (v1.11, parameter: k=15, -- MD) (Chirag et al., 2020) approach connects to the nextdenovo skeleton containing Gap, crosses the Gap and replaces sequences containing Gap regions to obtain a T2T level genome.

The location and number of genome gaps were used to determine the continuity of genomic assembly. The second-generation sequencing clean reads were compared to the reference genome using bwa (version: 0.7.17-r1188). The consistency of the assembled genome was evaluated by comparing the ratio and coverage of the second-generation sequencing. Then, BUSCO (version: 4.1.4; evalue=1e-05) was used to assess the genome’s completeness and accuracy.

### 2.4 Gene annotation

Repeat elements were identified using RepeatModeler (version: open-1.0.11, BuildDatabase -name mydb;RepeatModeler -database mydb -pa 10) (Smit et al. 2015) and LTR_FINDE (version: official release of LTR_FINDER_parallel, -threads 16, -harvest_out, -size 1000000, -time 300) to construct a denovo repeat sequence database. Based on denovo repetitive sequence libraries and the RepBase library (http://www.girinst.org/repbase) (Smit et al., 2015), using RepeatMasker (version: the open - 4.0.9, -nolow-no_is-norna-parallel 2) (Smith et al., 2013) repeated annotation;

RepeatProteinMask (version: open-4.0.9, -nolowsimplex -pvalue 0.0001) predicts the repeat sequence of TE_proteion type. All the repeats identified by different methods were combined into the final combined TEs after removing the redundant repeats.

Genome structure analysis was conducted using homology-based prediction, transcriptome-based prediction, and de novo prediction. For homology annotation, protein sequences of five closely related species, including *Aphis_gossypii, Acyrthosiphon_pisum, Bemisia_tabaci, Diaphorina_citri*, and *Diuraphis_noxia*, were downloaded from the National Center for Biotechnology Information database (NCBI) and used for annotation through genewise Exonerate (version: v2.4.0) (Slater and Birney, 2005). For the denovo prediction, Augustus (version: 3.3.2) and Genscan (version: 1.0) (Salzberg et al.,1998) were employed to annotate gene structures. For transcript annotation, the transcriptome data were aligned to the assembled genome using TransDecoder (version: v5.1.0), and nonredundant transcripts were obtained using Stringtie (version: 2.1.1). All the gene models were processed by MAKER (version: 2.31.10) (Holt and Yandell, 2011) to obtain the final gene set.

Gene function was annotated by searching the protein databases Uniprot (Apweiler et al., 2004), the Kyoto Encyclopedia of Genes and Genomes (KEGG) (http://www.genome.jp/kegg/) (Kanehisa and Goto, 2000), InterProScan (version: 5.52-86.0) (Blum et al., 2021), NR (Deng et al., 2006), KOG, and GO with blastp (version: 2.0.11.149, E-value <1e−5) (Buchfink et al., 2015). tRNAscan-SE (version: 1.23) (Chan et al., 2021) was used to annotate the noncoding RNA for tRNA screening and RNAmmer (version: 1.2) was used for sequence alignment with homologous species to identify rRNA. The Rfam database was used with the INFERNAL (version: 1.1.2) (Nawrocki and Eddy 2013) software to predict ncRNAs in the genome.

### 2.5 Phylogenetic analysis

A total of 16 insects were selected to construct a phylogenetic tree: *K. lacca, Ericerus pela, Phenacoccus solenopsis, B. tabaci, Nilaparvata lugens, Cimex lectularius, D. noxia, A. pisum, Drosophila melanogaster, Anopheles gambiae, Bombyx mori, Danaus plexippus, Tribolium castaneum, Anoplophora glabripennis, Nasonia vitripennis*, and *Daphnia pulex*. Protein sequences for these species were constructed with the orthofinder (version: 2.3.12, -M msa) pipeline using the default parameters.

We generated multiple sequence alignments for each 1:1 orthologous cluster using Muscle (Edgar, 2004). We used trimal (version: v1.4.rev22, -gt 0.2) to remove positions with gaps in more than 20% of the aligned sequences and concatenated all 1:1 orthologous genes in each species into a super-sequence. The phylogenetic tree was constructed using the maximum-likelihood method implemented in RAxML (version: 8.2.10) (Model: PROTGAMMAWAG) for branch support. Support values were obtained from 1,000 bootstrap replicates. Meanwhile, divergence time and age of fossil records were derived from TimeTree (http://www.timetree.org/) and applied as the calibration points. According to the divergence times from TimeTree, the nodal dates of *D. pulex* and *B. tabaci* were 452-557 million years ago (MYA), those of *E. pela* and *A. glabripennis* were 334-394 MYA, *N. lugens* and *A. pisum* were 177-401 MYA, *A. gambiae* and *D. plexippus* were 243-317 MYA, and those of *K. lacca* and *D. noxia* were 150.9-271.1 MYA.

Gene family expansion or contraction analysis was performed using CAFE (version: 4.2) (Han et al., 2013) based on the results of the orthofinder and the phylogenetic tree. The accepted level for indicating significantly altered gene families was p <0.05.

### 2.6 Positive selection analysis

We generated protein sequence alignments for single-copy gene families using MAFFT (localpair-maxiterate 1000). Then PAL2NAL (version: v14) was used to invert the protein sequence into a codon alignment sequence. The CodeML program from the PAML (version: 4.9i) package was implemented to estimate the dN/dS (nonsynonymous to synonymous) ratio for genes with the branch-specific model and the branch-site model. The likelihood ratio tests (LRTs) were used to compare the tested Model A (which allows positive selection on each of the foreground lineages, ω>1) with the null model, which does not allow such positive selection. The results of a significant difference (p < 0.05) were designated as “positively selected genes.”

### 2.7 HGT event identification

For *K. lacca*-specific HGTs, we submitted 10,696 predicted coding genes of *K. lacca* to a BLASTP search against the NCBI protein database (E-value=1×10^−5^). The proteins with the best BLAST hits in bacterial or fungal sequences were extracted, and the number of extracted sequences was counted to obtain the position and sequence information of HGT genes on the genome. After sequences without support from transcript evidence were excluded, a series of parameters were used to filter the candidates, including the sequences that were further compared to the NR library; the alignment consistency was 30-70%, the sequence length was >150 bp, and the q coverage was >50%. All candidate HGTs were subjected to phylogenetic analysis for verification. Synonymous codon-usage order values and GC contents of HGT and non-HGT genes were calculated by CodonO.

### 2.8 RNA sequencing and transcriptome assembly

Total RNA was extracted separately from L, A1, A2, A3, BH, BW, AT, and MG samples using the R6814 Blood RNA Kit (Life Technologies) following the procedure provided. RNA quantity, purity, and integrity were determined on a NanoPhotometer and an Agilent 2100 Bioanalyzer. Libraries were constructed using an Illumina NEBNext^®^ UltraTM RNA Library Prep Kit according to the manufacturer’s instructions and sequenced on an Illumina Novaseq 6000 platform. Clean reads were mapped to the genome assembly using Star (version: 2.7.9a, default) (Dobin et al., 2013). Differentially expressed genes were identified by padj <0.005 and log2 (fold change) >1. GO and KEGG enrichment was performed using clusterProfiler (version: 3.14.3) (Yu et al., 2012).

### 2.9 Predicting the putative lac biosynthesis pathway

Lac biosynthesis requires hydroxy-fatty acids and sesquiterpene esters, all of which participate in fatty acid biosynthesis and terpenoid biosynthesis. We used the KEGG PATHWAY and ORTHOLOGY tools in the Kyoto Encyclopedia of Genes and Genomes (KEGG) to detect all participating components and the KEGG orthology. Based on differential expression gene analysis in different developmental stages and different tissues, orthomcl version 2.0.9 with default parameters was used to detect orthologous groups of lac biosynthesis KOs in the sequenced *K. lacca* genome. blastp (E-value <1e-5) was used to determine the locations of the Kos. All steps related to lac biosynthesis were linked, and a KEGG pathway was proposed for lac biosynthesis.

*K. lacca* samples were prepared from different developmental stages (L, A1, A2, and A3) and different tissues (BH, BW, AT, and MG). Total RNA was extracted from the mixed sample of *K. lacca* using the EZ-10 Total RNA Mini-Preps Kit (Sangon Biotech, Shanghai, China) according to the manufacturer’s instructions. The first-strand cDNA was synthesized using the NovoScript Plus All-in-One 1st Strand cDNA Synthesis SuperMix (gDNA Purge) (Novoprotein Scientific Inc., Shanghai, China) according to the manufacturer’s protocol. Specific primers for the genes on the biosynthesis pathway of lac were designed using Primer Premier 6.0 (**Table S16**). In this study, the qPCR was performed using the NovoStart SYBR qPCR SuperMix Plus Kit (Novoprotein Scientific Inc.). The reactions were run on an ABI Quant Studio 3 real-time PCR system (Applied Biosystems, Thermo Fisher Scientific, Waltham, MA, USA). The gene expression levels were calculated with the 2^-ΔΔCT^ method (Livak and Schmittgen 2001).

## 3. Results

### 3.1 Genome assembly

We adopted Illumina and Oxford Nanopore Technologies sequencing for the comprehensive analysis of the *K. lacca* genome. A total of 68.06 GB of clean data was generated using the Illumina platform. The k-mer (K=19) analysis indicated that the heterozygosity of *K. lacca* was 0.27% and the estimated genome size was 262.92 Mb (**Table 1, Figure S1, Table S2**). The sequencing of the Oxford Nanopore Technologies, Oxford, UK platform generated 124.67 Gb raw data. After removing low-quality and short reads, we obtained 116.66 Gb of clean reads for genome assembly: a data set of 9,127,577 reads with an N50 of 34,324 bp (**Table S1**). A draft genome assembly of 258.04 Mb was obtained, yielding 26 contigs with a contig N50 of 23,848,510 bp.

**Table 1.** Statistics of the *K. lacca* genome assembly.

| Features | Statistics |
| --- | --- |
| Genome size (Mb) | 256,62 |
| Assembly level | Chromosome |
| Number of chromosomes | 9 |
| GC content (%) | 25.40% |
| Contig N50 (Mb) | 23,848,510 |
| Scaffold N50 (Mb) | 28,529,763 |
| BUSCO Insecta (%) | 95.7% |
| Protein-coding genes | 10,696 |

Then, we used Hi-C long-range scaffolding to improve our assembly. A total of 108.35 Gb of raw data was generated, and 92.73 Gb of filtered clean reads were used for the following analysis (**Table S1**). As a result, the chromosome-level genome has a total length of 256.62 Mb, with a scaffold of N50 of 28.53 Mb (**Table 1**). One hundred percent of the genome’s bases were successfully anchored to the nine chromosomes **(Figure S2, Table S3)**. Chromosome lengths ranged from 24,084,477 bp to 33,948,441 bp. Through T2T assembly technology, we finally obtained a high-quality genome with 0 gaps.

To assess assembly accuracy, we remapped the Illumina paired-end reads to the assembled genome. The reads mapped to 99.72% of the assembled genome sequences with a 99.23% mapping rate and 261.19 × average sequence depth (**Table S6**). Furthermore, 1,307 (95.7%) of the 1,367 highly conserved insect orthologues from BUSCO were identified in the assembly (**Table S5**). As revealed by BUSCO analyses against the Insecta datasets, the *K. lacca* genome assembly contained a higher number of conserved single-copies than the four published insect genomes (**Figure S3**). Together, these results suggest the completeness and high quality of our genome assembly.

We evaluated the genome of *K. lacca*. The survey result indicates that the proportion of commensal bacteria contamination was extremely high, and the effective data was only 22%. In order to remove the contamination of the lac insects, we adopted the strategy of increasing the sequencing volume, and the genome sequencing depth of the lac insects was 450 X, so as to ensure that the effective depth of assembly was 100 X. Nextdenovo’s assembly and comparison with NT showed that most of the sequences were aligned to the *K. lacca* genome. The GC_depth plot has no abnormal peak shape and conforms to the Poisson distribution, so the contamination cannot be investigated by the GC_depth plot alone (Figure S4). Then we used interaction signals in Hi-C to find the location of commensal contamination. The Hi-C-assisted assembly results show that the overall quality of our genome is high, but the last small piece of scaffold has a strong interaction signal of its own but no interaction with the rest of the genome, so we predicted that the last scaffold is contaminated with bacterial genes. To verify that it was a commensal contamination, NT alignment was performed on the last scaffold (named ctg000120_P) sequence with a length of 1,433,897 bp, and the results were mapped to the symbiotic bacteria *Wolbachia pipientis*. To further confirm the ctg000120_P contamination sequence for commensal bacteria that should be removed, we assembled the *Wolbachia pipientis* genome of size 1,539,149 bp. The 1,539,149 bp commensal sequences were compared with the 1,433,897 bp ctg000120_P with MiniMAP2, and 1,093,096 bp sequences of the 1,433,897 bp were found to be able to map the reassembled commensal sequences. Meanwhile, we used the assembled bacterial genome and the contig-level genome to compare with minimap2. Interesting, there were 521, 296 bp sequences that could be mapped to the reassembled symbiotic sequences, and their genome locations in lac insects were all chimeric within contigs with an extremely low proportion, which suggests that ctg000120_P may be caused by horizontal gene transfer (HGT) rather than bacterial contamination. In conclusion, the bacterial sequence found in the genome is the result of HGT and pollution, with a 32.29% chance that it was generated by HGT and a 67.71% chance that it was caused by pollution.

### 3.2 Genome annotation

A total of 79,136,004 bp of repetitive sequences were obtained in the *K. lacca* genome, yielding a repeat percentage of 22.19% **(Table S7)**. *K. lacca* had far fewer repetitive sequences than the closely related species *Ericerus pela* (37.3%) and *Phenacoccus solenopsis* (37.9%) (Chen et al., 2021; Li et al., 2020). We integrated expression data from eight RNA-seq libraries, homolog protein alignment, and de novo gene prediction to create 10,696 protein-coding genes in *K. lacca*. The average CDS length, exon number per gene, exon length, and intron length were 1,342.95 bp, 6.97 bp, 269.31 bp, and 1,321.07 bp, respectively, similar to those in most of the reported scale insect species **(Table S8)**. This BUSCO analysis indicated that the annotated gene-sets have a completeness value of 94.2% (**Table S5**). Among the 10,696 predicted genes, 9,599 (89.74%) were functionally annotated, including 9,199 (86.00%) genes found via the Interproscan database, 8,850 (82.74%), and 8,839 (82.64%) genes via the NR and Uniprot databases, respectively (**Table S9**).

### 3.3 Gene orthologues and comparative genomic analysis

We used the OrthoMCL to identify orthologous genes among *K. lacca* and 15 other insect species. A total of 37,306 gene family clusters were constructed, with 446 identified as 1:1:1 single-copy orthologous genes (**Table S12**). In addition, we also identified 880 *K. lacca* specific gene families. Using the protein-coding sequences of 446 single-copy orthologous genes (*Daphnia pulex* as an outgroup), a phylogenetic tree was constructed. The results showed that *K. lacca* and seven other Hemiptera were clustered together. Scale insects (including *K. lacca, E. pela*, and *D. pulex*) and Aphididae (including *Diuraphis noxia* and *Acyrthosiphon pisum*) diverged from their common ancestor about 169.4 million years ago (MYA) (156.5∼178.5 MYA). *K. lacca* was a sister taxon to *E. pela*. The two scale insect species diverged from their common ancestor at approximately 57.9 MYA (44.6∼70.3 MYA). Further, our analysis indicated that the last common ancestor of holometabola and incomplete metamorphosis separated in the Late Devonian 389.5 (372.6∼409.3 MYA), at least 80 million years earlier than the previous study (Misof et al., 2014; Chen et al., 2021). Based on this phylogenetic analysis, ametabolous insects represent a primitive group located in the basal branch of the tree, whereas Hemiptera and holometabola reside separately in the lower and upper branches. Three scale insects (*K. lacca, E. pela*, and *D. pulex*) are an important transitional taxon between incomplete metamorphosis and holometabola. This similarity to a previous report supports an evolutionary trend from ametabolous to incomplete metamorphosis insects, followed by the emergence of holometabola (Chen et al., 2021) (Figure 2-B).

**Figure 1.**
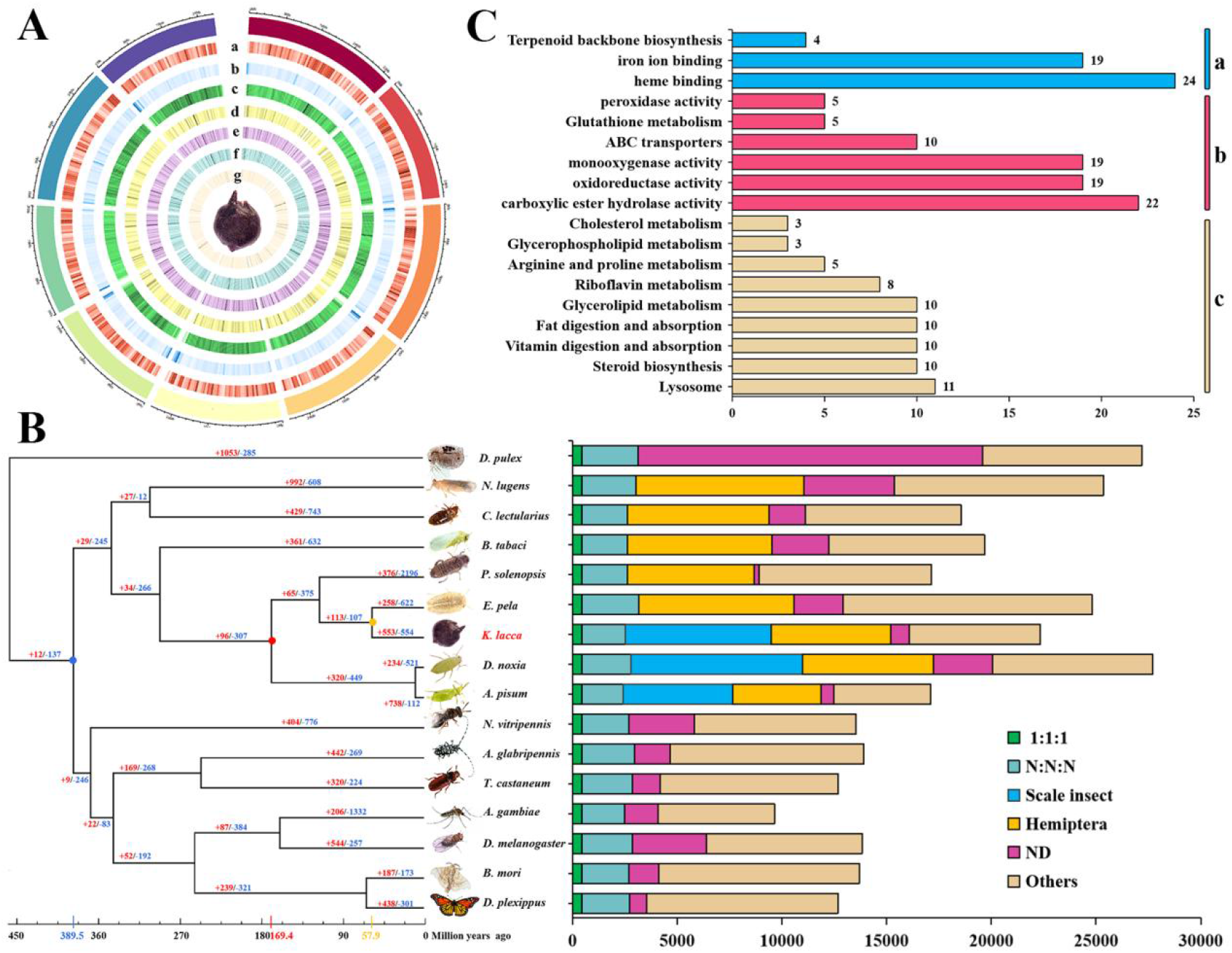
Genome evolution of *kerria lacca*. **A**. Distribution of *K. lacca* genomic features. Track a-c: gene density (50 kb window), repeats density (50 kb window), and GC content (50 kb window). Track d-g: Mean expression of annotated genes calculated as lg^FPKM^ of the mean expression value in different developmental stages (A1, A2, A3, and L). **B**. Phylogenetic analysis of *K. lacca* with other insect species. The phylogenetic position of *K. lacca* was determined based on 446 single-copy genes. The estimated species divergence time is illustrated at the bottom of the phylogenetic tree. The gene family expansion (red) and contraction (blue) are illustrated at the branches and nodes of the tree. The blue dot and number indicate the divergence time of the incomplete metamorphosis separated from the holometabolan, red color show the time of scale insect diverge from aphids and those in orange color indicate the time of *K. lacca* diverge from *E. pela*. 1:1:1 indicates single copy genes, and N:N:N indicates multicopy genes across 16 insect species. The Scale insects and Hemiptera indicate suborder-specific genes, respectively. **C**. Enrichment analysis of expanded genes of *K. lacca*. a. *K. lacca*-specific related genes. b. Detoxification genes. c. Nutritional metabolism-related genes.

**Figure 2.**
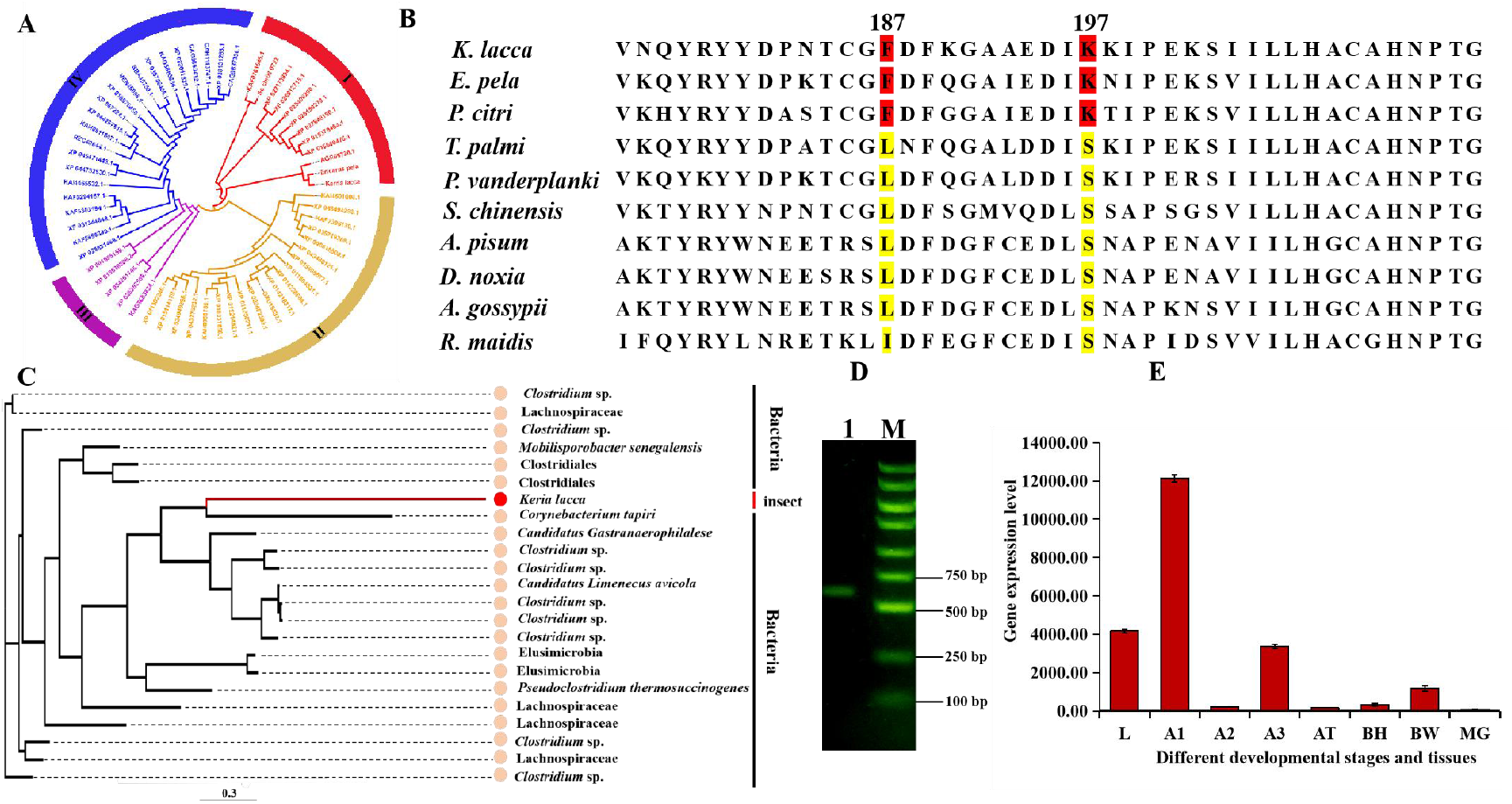
Positive selection and Horizontal gene transfer. **A**. Phylogenetic tree of the AST gene in *K. lacca* and 58 other insects. **B**. Alignment of three scale insects and seven other insects AST amino acid sequences. Amino acids unique to the three scale insect species (corresponding to residues 187 and 197 in other insects AST) are shown in red; other amino acids at these positions are shown in yellow. **C**. Maximum likelihood phylogenetic analysis of IPPS. **D**. Genome fragments cloned from *K. lacca* IPPS. M, DNA marker; 1, PCR products of the IPPS gene fragment. **E**. Gene expression levels of IPPS in different development stages and tissues.

Significant expansion and contraction of gene families are usually related to the adaptive divergence of species. Three scale insects showed 375 expanded and 65 contracted gene families compared with those of the common ancestor of (*K. lacca, E. pela*, and *D. pulex*) and two Aphididae (*Diuraphis noxia* and *Acyrthosiphon pisum*), while the *K. lacca* genome displayed 258 expanded and 622 contracted gene families compared with those of the common ancestor **(Figure 2-B)**. We analyzed the functional properties of the expanded 258 gene family (758 genes), and KEGG and GO enrichment analysis revealed that the identified 70 gene expansion might be related to enriching the nutrient metabolism pathway of the lac insect. The expanded gene families were involved in Lysosome (11), vitamin digestion and absorption (10), fat digestion and absorption (10), glycerolipid metabolism, and so on. Because of its piercing sucking mouthpiece, the lac insect can only absorb a limited amount of nutrients from its host plants. The expansion of nutrition metabolism genes might facilitate the digestion and absorption of nutrients from plant hosts. In addition, 80 genes were significantly enriched in categories primarily involved in xenobiotic detoxification, such as carboxylic ester hydrolase activity (22), oxidoreductase activity (19), monooxygenase activity (19), ABC transporters (10), and so on. The expansion of genes might facilitate the detoxification of natural xenobiotics from plants (Figure 2-C, **Figure S6-B**).

Three groups of 47 genes involved in the *K. lacca*-specific related gene also underwent expansion. Among them, four genes expanded that were functionally related to terpenoid biosynthesis. This might be because lac insect species produce a large amount of lac, which is an effective adaptation for *K. lacca* to escape predators and cope with harsh environmental conditions. In addition, 24 genes involved in heme binding and 24 genes involved in an iron ion binding-related gene also underwent expansion; this might be related to the red pigment synthesis of the lac insect (**Figure** 2-C, **Figure S6-A**).

### 3.4 Positive selection analysis

To further uncover the genomic characteristics of *K. lacca*, a positive selection analysis was performed. Using 446 single-copy gene families, 49 positive selection genes (PSGs) (FDR < 0.05) were identified. GO and KEGG enrichment analysis, we found that 11 PSGs participated in pathways that may be associated with nutrition metabolism, such as the amino acid metabolism (aspartate aminotransferase (AST)), glycerophospholipid metabolism (phospholipase), and glycan biosynthesis (glucoside xylosyltransferase). Moreover, PSGs were also related to the cell cycle, RNA transport, and ribosome formation. Amino acids in insect haemolymph play a key role in providing energy through osmosis, proline metabolism, neurotransmission, and other metabolic pathways (Klowden, 2007; Mirhaghparast et al., 2015). There are reports that AST is a sign of the entrance of amino acids into the glucogenesis process (Lehninger, 1982). Gluconeogenesis is the main pathway for sugar synthesis from non-carbohydrate substrates (Etebari et al., 2007). In this series of reactions, the carbon source of gluconeogenesis is amino acids, and the activation of this metabolic pathway is usually related to the large reduction of free amino acids in fat bodies and hemolymph. At this time, alanine is converted into pyruvate to provide energy (Matindoost et al., 2005; Hasheminia et al., 2011). We analyzed the aspartate aminotransferase (AST) gene, which is a positive selection in the genome. It was composed of 428 amino acids. On the basis of 59 AST protein sequences from three scale insects and other insects, we constructed a phylogenetic tree. The gene phylogeny strongly supported the interspecific relationships demonstrated in a previous tree based on species classification analyses (Figure 2-A). In the analysis of the AST amino acid sequence of *K. lacca*, it was found that it contained multiple conserved domains, including the unique conserved domain of the aspartate aminotransferase supergene family. We then identified shared amino acid substitutions that occurred along the lineage containing scale insects *K. lacca, E. pela*, and *D. pulex*, but not along the lineages leading to the Hemiptera (*Acyrthosiphon pisum, Aphis gossypii, Diuraphis noxia, Rhopalosiphum maidis, Schlechtendalia chinensis*), Thysanoptera (*Thrips palmi*), and Diptera (*Polypedilum vanderplanki*) species. We speculate that amino acid substitution in the conserved domain of the three scale insects might contribute to energy conversion and thus provide energy (Figure 2-B).

### 3.5 Horizontal gene transfer

Horizontal gene transfer is a significant dynamic of genome evolution in both prokaryotic and eukaryotic organisms, and it is accompanied by adaptive evolution (Li et al., 2022). We used a variety of methods to identify the genes that may cause HGT in *K. lacca*. First, 27 candidate genes were compared to fungi or bacteria through protein-coding genes of the genome and compared to the NR library by blast. Then, 16 candidate genes were obtained through screening genes supported by transcripts. Finally, the above sequences were compared to the NR library, where the alignment consistency was 30-70%, the sequence length was >150 bp, and the q coverage was greater than 50%, yielding 5 genes that underwent HGT events.

Candidate gene *Kerria0046210*.*1* encodes Isoprenyl diphosphate synthases (IPPS) through achieving comparison in the NR library. The phylogenetic tree of Isoprenyl diphosphate synthases (IPPS) of 23 species was constructed by blast and RAXML. The result that the gene from *K. lacca* and *Corynebacterium tapiri* formed a branch and had a close affinity shows the gene may undergo horizontal gene transfer (Figure 2-C). The GC content calculated by CodonO of the gene is higher than that of adjacent upstream and downstream genes (**Table S13**). Transcriptome analysis in different developmental stages and tissues showed the gene has high expression in each stage and tissue of *K. lacca*. The PCR results showed that the *Kerria 0046210*.*1* gene existed in *K. lacca* (Figure 2-D). The protein encoded by the gene is composed of 242 amino acids. The analysis of the amino acid sequence of *Kerria 0046210*.*1* showed that it had a conserved domain of the cis-isopentenyl transferase family using the NCBI’s Sequence Conservative Domains (CD) search tool. Previous studies have shown that IPPS is a key enzyme connecting the biosynthetic branching points of upstream mevalonate and downstream isoprene (Wang and Ohnuma, 2000; Chen et al., 2021). The back of the body wall (BW), anal tubercle (AT), and the brachia (BH) process with more glands have higher expression compared with the midgut without glands in different tissues through differential gene analysis (Figure 2-E). Therefore, isopentenyltransferase plays an important role in the process of terpenoid skeleton biosynthesis and regulation, and it may be the key gene for lac synthesis.

### 3.6 The putative lac biosynthesis pathway

To construct a putative lac biosynthetic pathway in *K. lacca*, two related KEGG pathways were analyzed: fatty acid synthesis and terpenoid biosynthesis. Comparative genomic analysis of *K. lacca* showed that 3 of the genes that were related to fatty acid biosynthesis and 4 genes involved in terpenoid biosynthesis underwent expansion in the expansion gene family. Furthermore, we used transcriptome analysis to screen 25 key candidate genes in the lac biosynthesis pathway in different developmental stages and tissues. The different developmental stages that the lac secretion-active adult stages have in comparison with the lac secretion-minimum larval stage, in the process of lac biosynthesis, 11 genes were differentially expressed. Different tissues with more glands, BH, BW, and AT, had 14 genes with differential expression when compared to glandless MG. There were 10 candidate genes related to terpenoid biosynthesis and 15 related to fatty acid biosynthesis. The q-PCR results of the above key genes were basically consistent with the expression profile found with RNA-seq (Figures 3-A and 3-B). Based on these analyses, we proposed a putative pathway for lac biosynthesis. Though further validation is required, this putative pathway integrates most steps of lac biosynthesis, which is a useful resource for elucidating the biosynthesis mechanism of lac (Figure 3-C).

**Figure 3.**
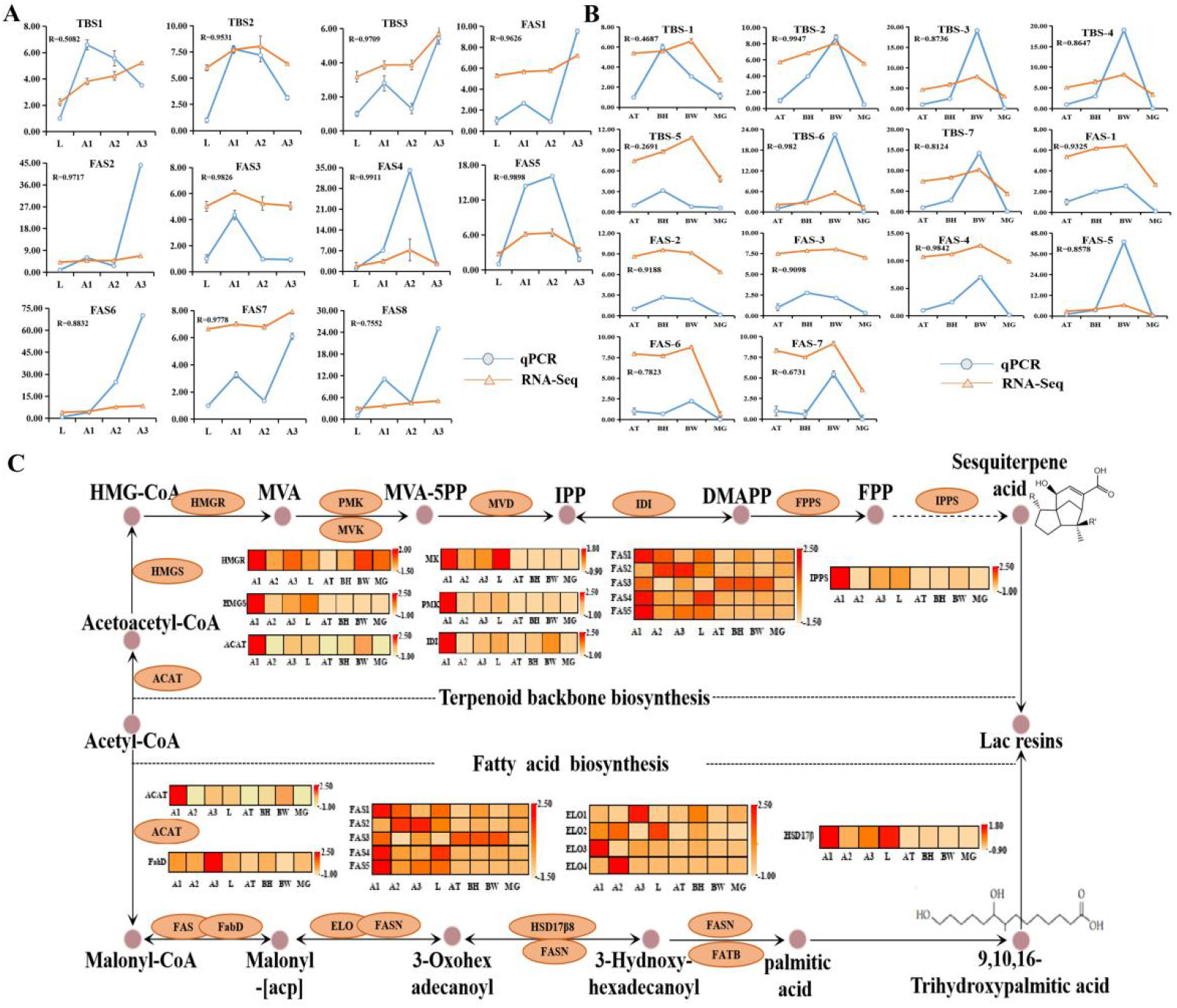
Comparative transcriptomic analysis of genes involved in the lac biosynthetic pathway. **A**. Genes expressed differentially at different developmental stages. **B**. Genes expressed differentially at different tissues. The horizontal represents different developmental stages and tissues of the lac insect. The numbers on the vertical axis represent gene expression for the respective genes. **C**. Comparative transcriptomic analysis of genes involved in the lac biosynthetic pathway of *K. lacca* from different developmental stages and different tissues.

## 4 Discussion

Scale insects contain a large number of endosymbiotic bacteria (Baumann, 2005; Szabó et al., 2017; Kandasamy et al., 2022), which poses a great challenge to getting high-quality genomes. In this study, we increased the amount of sequencing and then assembled the genome at the contig level for GC_ depth map, NT library pair, and interaction signal in Hi-C to obtain a high-quality genome from lac insect contaminated by a large number of symbiotic bacteria as well as a portion of horizontally transferred genes. In order to get rid of symbionts contamination and get horizontally transmitted genes, this will provide a new strategy for subsequent sequencing of scale insects and their highly contaminating species.

Horizontal gene transfer (HGT) is the transfer of genetic information from one organism to another via indirect inheritance in prokaryotes, phagocytic cells, and parasitic unicellular eukaryotes (Gladyshev et al., 2008). Insects inherit most of their horizontally transferred genes from bacteria and fungi, followed by a few viruses and plants. HGT is accompanied by the adaptive evolution of insects and plays an important ecological role in the functional adaptation of insects. For example, the carotenoid biosynthesis gene was transferred from fungi to aphids, which can change the body color of aphids (Moran and Jarvik, 2010); the whitefly detoxifying ability was enhanced by phenolic glycoside genes obtained from plants (Xia et al., 2021); the parasitoid killing factor (PKF) gene was introduced to Lepidoptera from a virus to increase their defenses (Gasmi et al., 2021); and the LOC105383139 gene, which was transferred from a bacterium to *Plutella xylostella* to enhance male-female courting behavior (Li et al., 2022). Scale insects include a large number of endosymbionts that are important in their nutrition and metabolism (McCutcheon and Moran, 2012; Husnik et al., 2013). These symbionts play key roles in the biology of their host insects, providing nutrients like essential amino acids and vitamins that the insects are unable to produce (Baumann, 2005; Douglas, 1989; Moran, 2007). During the long-term evolution of symbiotic bacteria and insects, several genes from symbiotic bacteria were horizontally transferred to the insect genome and carried out their biological functions. We found an IPPS gene that had been horizontally transferred from bacteria into the scale insect. Studies have shown that isopentenyl transferase (IPPS), also known as *Hevea* rubber transferase (HRT), is found in the final step of natural rubber biosynthesis and is a key enzyme in rubber biosynthesis. Its family can interact with farnesyl pyrophosphate (FPP), SRPP, HRBP, and REF to regulate the activity of these essential proteins, and so it plays an important role in the regulation of rubber biosynthesis (Lou et al., 2009; Yamashita et al., 2016; Tang et al., 2016). In the rubber biosynthesis of *Hevea brasiliensis*, rubber particles (RPs) are special organelles for rubber biosynthesis and storage, and the most abundant proteins on the RPs membrane are IPP, Z-IPPS, small rubber particle protein (SRPP), and rubber elongation factor (REF), of which Z-IPPS is a significant large component that impacts not only the speed of rubber biosynthesis but also the production of latex (Wang et al., 2010; Liu et al., 2020). Researchers found that IPPS is involved in rubber synthesis in the rubber-producing plant (*Taraxacum koksaghyz*), and its activity is positively correlated with rubber production (Schmidt et al., 2010). Eight IPPS genes were identified in its genome, and one of them, IPPS1, is highly expressed in latex, providing more target genes for subsequent specific functional studies of IPPS genes (Li et al., 2017; Xie et al., 2019). Unlike the rubber tree, which produces *cis*-polyisoprene, *Eucommia ulmoides* rubber biosynthesis depends on the *trans*-isopentenyl transferase E-IPPS, in which the farnesyl diphosphate synthases (FPS) plays a role in the synthesis of natural trans rubber as E-IPPS (Wuyun et al., 2017). Therefore, the IPPS gene, which was horizontally transferred from bacteria into the scale insect genome, is a key enzyme that connects the biosynthetic sites of upstream mevalonate and downstream isoprene, plays an important role in the biosynthesis and regulation of terpenoid backbones, and might be a key gene of lac synthesis. In-depth investigation of the specific biological function of IPPS in the regulation of lac biosynthesis in lac insects will lead to new insights for elucidating the precise mechanism of lac biosynthesis and secretion.

Based on Illumina and ONT sequencing platforms, Hi-C technology, and T2T assembly, we provide the genome of *K. lacca* at chromosome level 0 Gap. The phylogeny suggests that *K. lacca* diverged from a common ancestor with *E. pela* 57.94 MYA. Among the expanded 258 gene families, 197 genes related to nutrition metabolism, detoxification and *K. lacca*-specific were identified, and positive selection occurred in AST genes related to nutrient metabolism in positive selection analysis. By transcriptome analysis of different developmental stages and tissues, 25 candidate genes related with *K. lacca* synthesis were screened, among which IPPS genes from bacteria were integrated into the *K. lacca* genome via horizontal gene transfer, and the biosynthetic pathway of *K. lacca* was constructed. This study will provide a basis for the analysis of the unique nutritional metabolism mechanism, as well as genetic and evolutionary studies of the scale insect.

## Acknowledgements

The research was supported by the grant for Fundamental Research Funds of CAF (CAFYBB2020QA003), National Natural Science Foundation of China (31772542) and grants for the high level foreign experts introduced to Yunnan province (YNQR-GDWG-2019-011).

## Author contributions

X.M.C. and H.C. designed and led the project. W.W.W. and N.H.B. collected samples. W.W.W., Q.L. and W.F.D. accomplished the whole-genome sequencing and Hi-C assembly. W.W.W., H.C., Q.L., and W.F.D. performed whole-genome assembly quality control and transcriptome analyses. W.W.W., J.W.Z. and X.F.L afforded photos of fresh collection and designed the figures; W.W.W., N.H.B., Q.L. and H.C. wrote the manuscript. X.M.C. and H.C. cowrote and edited the manuscript. All authors read and approved the final manuscript.

## Data availability statement

The datasets generated and analysed during the current study are available in the Sequence Read Archive repository (SRA; https://www.ncbi.nlm.nih.gov/sra/). The RNA-seq data generated in this study have been submitted to the NCBI BioProject database (https://www.ncbi.nlm.nih.gov/sra/PRJNA896359) under accession number PRJNA896359. Hi-C data (the chromosome-level assembly) have been submitted to the Sequence Read Archive (SRA; http://www.ncbi.nlm.nih.gov/bioproject/922917) under accession number PRJNA922917. The final DNA sequence assembly has been deposited at DDBJ/ENA/GenBank under the accession LAC00000000.

