## Supplementary Information for "High-resolution lac insect genome assembly provides genetic insights into lac synthesis and evolution of scale insects"

### 1.Supplementary Figures

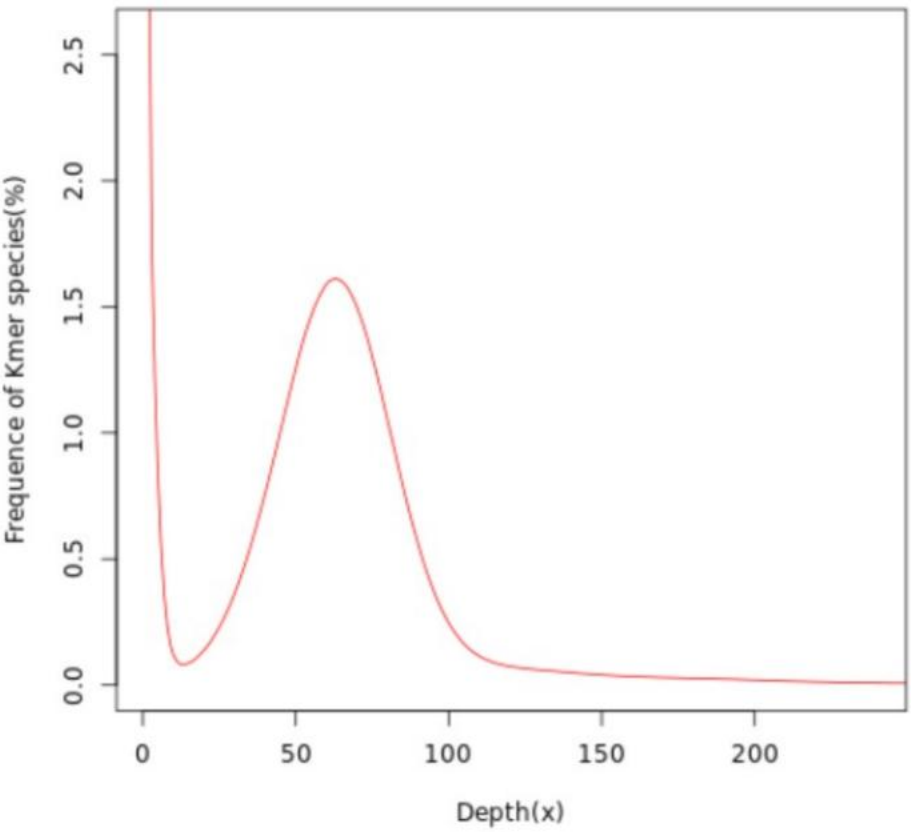


**Figure S1 Distribution of 19-mer frequency in *Kerria lacca* genome.**

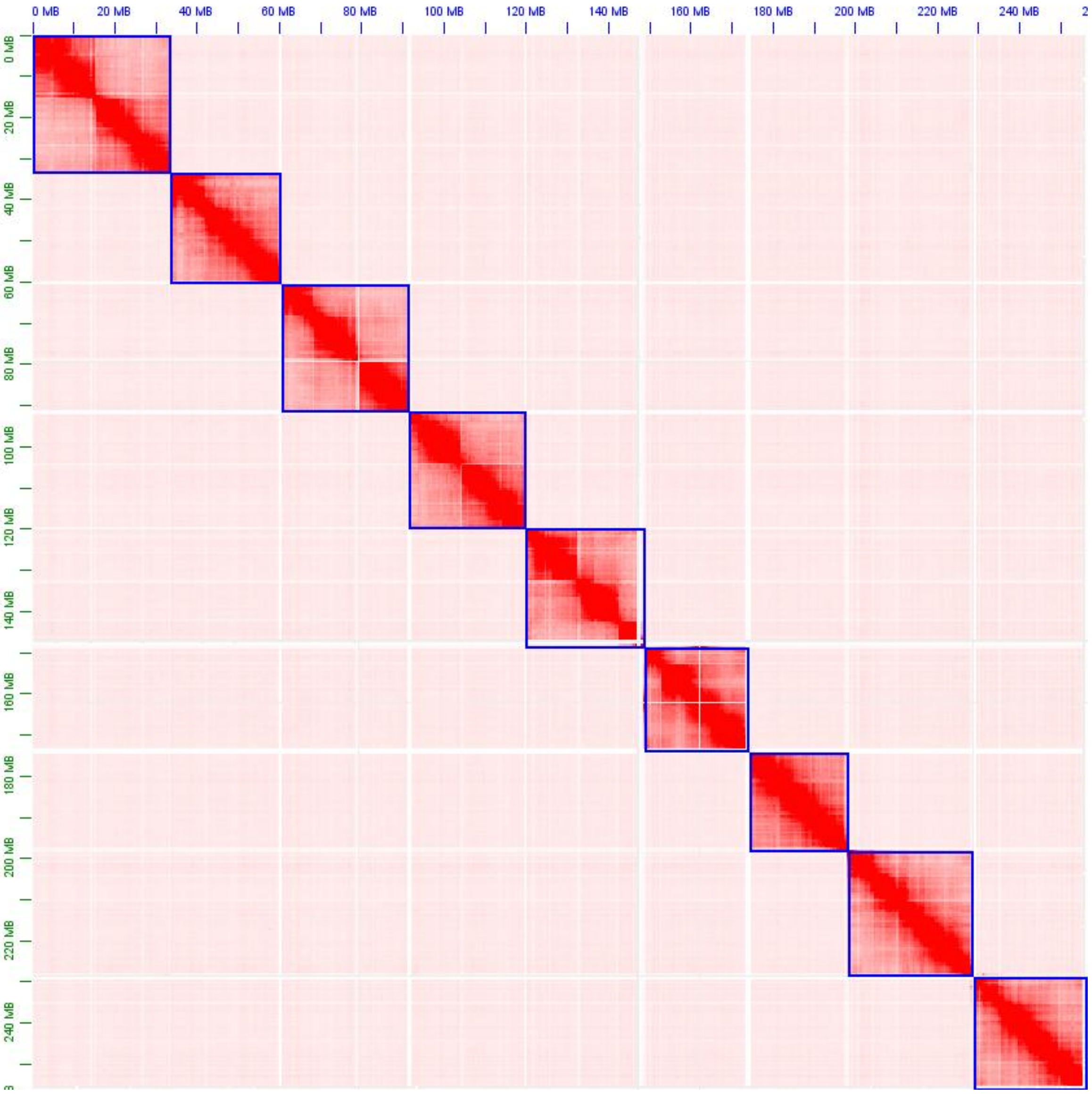


**Figure S2 Contact maps of Hi-C interactions among chromosomes in the *K. lacca* genome.**

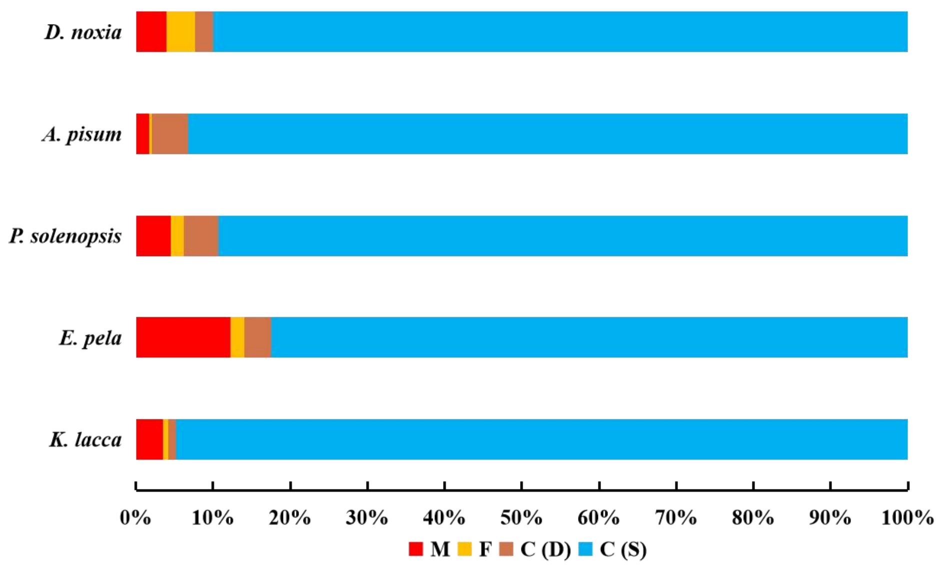


**Figure S3 Assessments of BUSCO completeness. C:** complete BUSCOs, **S:** complete and single-copy

BUSCOs, **D:** complete and duplicated BUSCOs, **F:** fragmented BUSCOs, **M:** missing BUSCOs.

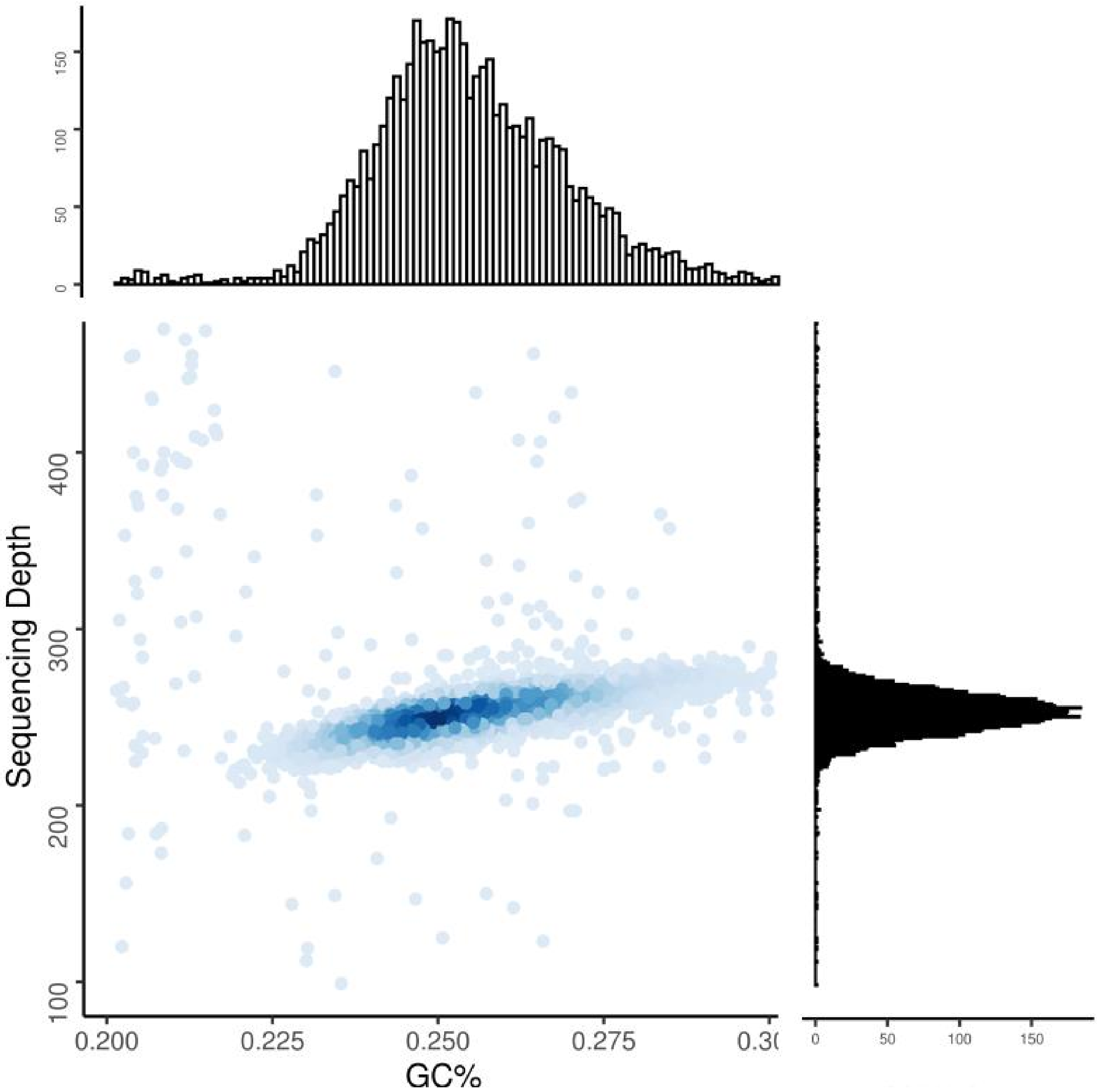


**Figure S4　GC content against the sequencing depths of *K. lacca* genome.** The x-axis represents the GC content, and the y-axis represents the depth of sequencing. The histogram (up) represents distribution of GC content, and the histogram (right) represents distribution of sequencing depth.

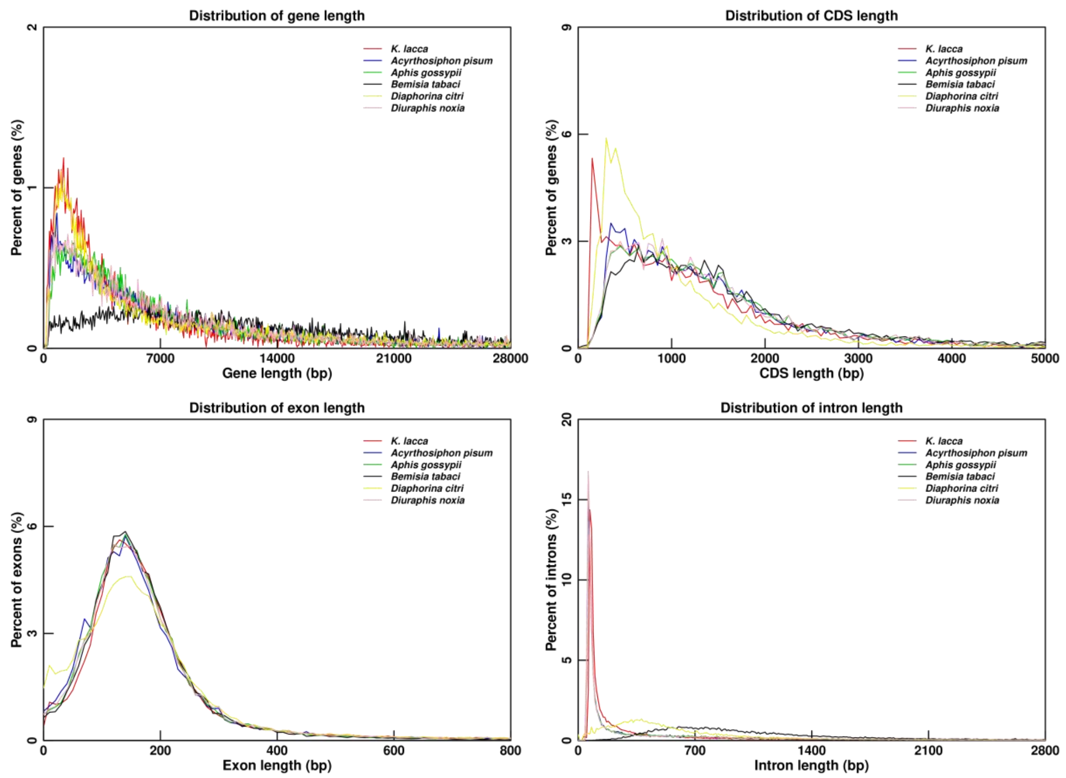


**Figure S5  Characteristics of the annotated protein-coding genes in the *K. lacca* genome**

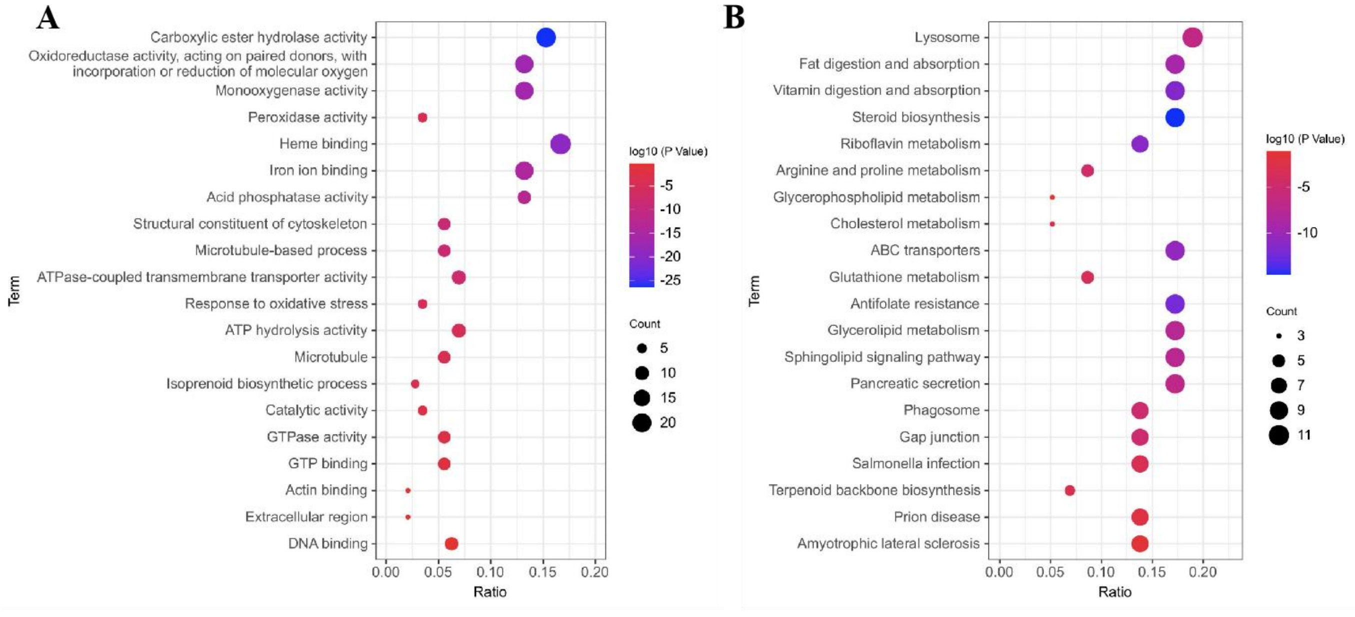


**Figure S6  enrichment analysis of expanded gene families of *K. lacca*. A**. Gene ontology (GO) enrichment analysis of expanded gene families of *K. lacca*. **B.** KEGG pathway enrichment analysis was performed for the expansion gene family of *K. lacca*.

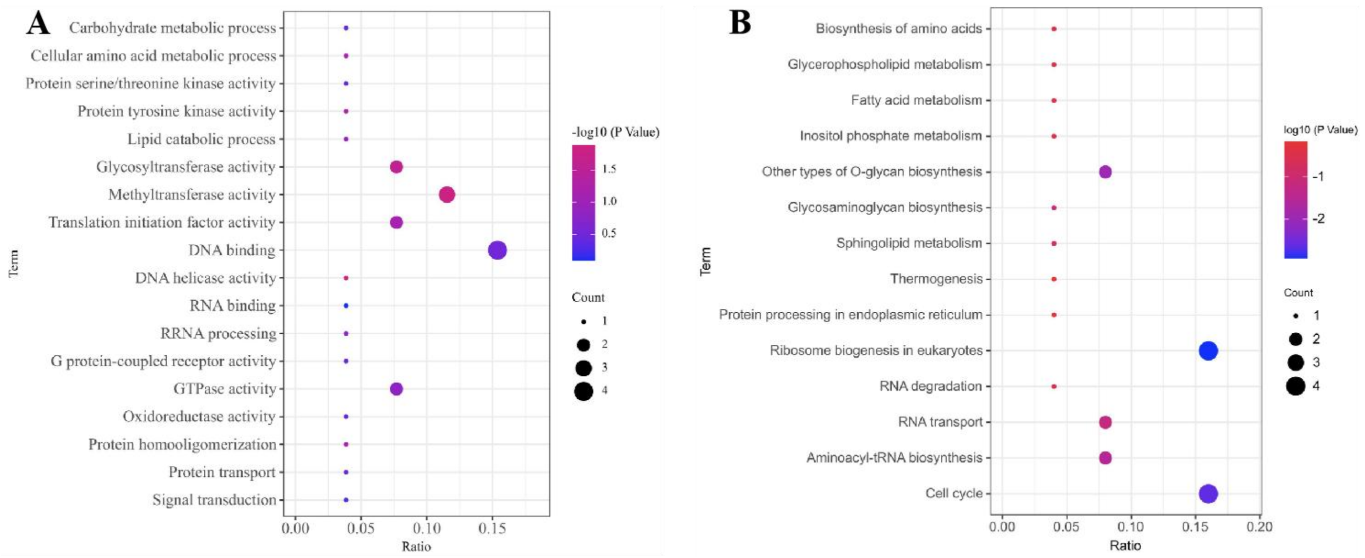


**Figure S7 enrichment analysis of Positive selection of *K. lacca*. A.** Gene ontology (GO) enrichment analysis of positive selection of *K. lacca*. **B.** KEGG pathway enrichment analysis was performed for the positive selection of *K. lacca*.

```
1    ATGAAAGAATTTGCAAAAAAAATTTTTATTCCGATTTTAACGTTT
      M  K  E  F  A  K   K  I  F  I  P  I   L  T  F
46   ATCGAAAATCTTTTATTTAAATTATGCGTTTTCGTTATAAAGCAG
      I  E  N  L  L  F  K  L  C  V  F  V  I  K  Q
91   AATGCTGTACCAAAACACATTGCACTTATCGTGGATGGGAATCGA
      N  A  V  P  K  H  I  A  L  I  V  D  G  N  R
136  TCATTTTCAAAATCGAGCGGAGTTTCATTAGATGATGCATATATG
      S  F  S  K  S  S  G  V  S  L  D  D  A  Y  M
181  AAAGGCGTATTTAATTGGGACAGAAAAAAATCAGAAACTGAACCT
      K  G  V  F  N  W  D  R  K  K  S  E  T  E  P
226  ATTTTGAAGGCGATTACAGCTCTCATGAAGATGTCTATGGCTAAT
      I  L  K  A  I  T  A  L  M  K  M  S  M  A  N
271  ACATACGGATGTCGATACAAATTAGTCGGTGACATCTCTAAATTG
      T  Y  G  C  R  Y  K  L  V  G  D  I  S  K  L
316  CCGCCTGATTTTTTAGCATTAGGTAAAGATTTAGAAAACAAGACT
      P  P  D  F  L  A  L  G  K  D  L  E  N  K  T
361  AAATATAATACCGGAGCGGTTATTCAATTAGCTGTAAATTACGGA
      K  Y  N  T  G  A  V  I  Q  L  A  V  N  Y  G
406  GCTAAATTAGAAATGGTCGATTCTGCGTTAGCAATTTTCAAACAT
      A  K  L  E  M  V  D  S  A  L  A  I  F  K  H
451  CCTTCATATTTACAAAATATAACCGAAGAACTAATCGAAGATTTG
      P  S  Y  L  Q  N  I  T  E  E  L  I  E  D  L
496  ATGAATGTCCACGGAGTTGAAAAAACAGATATAATGATTCGAACT
      M  N  V  H  G  V  E  K  T  D  I  M  I  R  T
541  TCCGGACATAAAAGATACAGCGATTATTTTTTGTGGCAAAACGCT
      S  G  H  K  R  Y  S  D  Y  F  L  W  Q  N  A
586  TACCATCCTTTATTCTTTACCGATGAATTTTGGCCTGAATTTTCA
      Y  H  P  L  F  F  T  D  E  F  W  P  E  F  S
631  CGTTGGGGTTTGTTAAGAGCGGTTTTTTATTATCAAGCTAATATA
      R  W  G  L  L  R  A  V  F  Y  Y  Q  A  N  I
676  AAAAAATGTTTAAGAGCAAAAGCGATTGCTCGGAGAAATACTTTT
      K  K  C  L  R A  K  A  I  A  R  R  N  T  F
721  AAATAA
      K  *
```

**Figure S8 Nucleotide and deduced amino acid sequence of the *Kerria 0046210.1* gene.**

### 2.Supplementary Tables

**Table S1 The statistics of sequencing data used for the *Kerria lacca* genome assembly**

| Libraries | Insert size (bp) | Raw data (Gb) | Clean data (Gb) | Clean reads (Gb) | Sequence coverage (fold) |
|---|---|---|---|---|---|
| Illumina reads | 150 | 69.00 | 68,06 | 457,671,624 | 300 |
| PacBio reads | 15 k | 124.67 | 116.66 | 9,127,577 | 450 |
| Hi-C reads | -- | 108.35 | 92.73 | 618,171,038 | 400 |

**Table S2 Estimation of *K. lacca* genome size using K-mer analysis**

| K-mer | K-mer number | K-mer Depth | Genome Size (Mb) | Revised Genome Size* (Mb) | Data Size (G) | X | Heterozygous Retio (%) | Duplication Ratio(%) |
|---|---|---|---|---|---|---|---|---|
| 19 | 16,777,202,489 | 63.81 | 262.92 | 258.04 | 27.07 | 102.96 | 0.27 | 22.11 |

* Revised genome size is accurate estimation without error rate.

**Table S3 Estimation of *K. lacca* genome assembly**

| | Qv | completeness(%) | length | N50 | Reads Number |
|---|---|---|---|---|---|
| initial genome | 35.6781 | 98.5498 | 256,439,521 | 23,705,252 | 26 |
| Finally genome | 41.9091 | 98.7307 | 256,623,901 | 28,534,742 | 9 |

**Table S4 T2T assembly of the *K. acca* genome**

| Chromosome | Length(bp) | contig number | gap number | QV |
|---|---|---|---|---|
| chr1 | 33,948,441 | 1 | 0 | 43.60 |
| chr2 | 26,625,331 | 1 | 0 | 45.70 |
| chr3 | 31,300,509 | 1 | 0 | 41.46 |
| chr4 | 28,529,763 | 1 | 0 | 42.37 |
| chr5 | 28,735,584 | 1 | 0 | 40.16 |
| chr6 | 25,618,100 | 1 | 0 | 40.91 |
| chr7 | 24,084,477 | 1 | 0 | 43.49 |
| chr8 | 30,300,621 | 1 | 0 | 43.80 |
| chr9 | 27,461,952 | 1 | 0 | 39.26 |

**Table S5 BUSCO evaluation of the genome assembly and annotation**

| Item | Assembly | | Annotation | |
|---|---|---|---|---|
| | Proteins | Percent(%) | Proteins | Percent(%) |
| Complete BUSCOs | 1,307 | 95.7 | 1,288 | 94.2 |
| Complete Single-Copy BUSCOs | 1,294 | 94.7 | 1,261 | 92.2 |
| Complete Duplicated BUSCOs | 13 | 1.0 | 27 | 2.0 |

| | | | | |
|---|---|---|---|---|
| Fragmented BUSCOs | 11 | 0.8 | 10 | 0.7 |
| Missing BUSCOs | 49 | 3.5 | 69 | 5.1 |
| Total BUSCO groups searched | 1,367 | 100.0 | 1,367 | 100.0 |

**Table S6** The alignment information of reads mapping to the *K. lacca* genome

| Reads | Mapping rate (%) | 99.23 |
|---|---|---|
| Genome | Paired mapping rate(%) | 97.38 |
| | Average sequencing depth | 261.19 |
| | Coverage (%) | 99.72 |
| | Coverage at least 4X (%) | 99.46 |
| | Coverage at least 10X (%) | 99.25 |
| | Coverage at least 20X (%) | 99.06 |

**Table S7** **The ratio of TEs in *K. lacca* genome.**

| Type | TE protiens | | Denovo+Repbase | | Combined TEs | |
|---|---|---|---|---|---|---|
| | Length (bp) | % in Genome | Length (bp) | % in Genome | Length (bp) | % in Genome |
| DNA | 3,088,312 | 1.20 | 6,226,227 | 2.43 | 8,210,831 | 3.20 |
| LINE | 4,513,197 | 1.76 | 9,584,184 | 3.73 | 11,023,402 | 4.30 |
| SINE | 0 | 0.00 | 104,693 | 0.04 | 104,693 | 0.04 |
| LTR | 1,168,250 | 0.46 | 2,001,616 | 0.78 | 2,436,187 | 0.95 |
| LTR-Gypsy | 570,692 | 0.22 | 1,352,914 | 0.53 | 1,510,446 | 0.59 |
| LTR-Copia | 256,901 | 0.10 | 264,586 | 0.10 | 374,121 | 0.15 |
| Satellite | 0 | 0.00 | 25,159 | 0.01 | 25,159 | 0.01 |
| Simple_repeat | 0 | 0.00 | 0 | 0.00 | 0 | 0.00 |
| Other | 0 | 0.00 | 12,944 | 0.01 | 12,944 | 0.01 |
| Unknown | 6,621 | 0.00 | 30,776,120 | 11.99 | 30,782,741 | 12.00 |
| Total | 8,773,441 | 3.42 | 47,928,458 | 18.68 | 56,935,999 | 22.19 |

**Table S8** **Genome annotation of *K. lacca***

| Item | Number |
|---|---|
| the total number of gene | 10,696 |
| the average of mRNA_length (bp) | 9,769.37 |
| the average cds_length of per gene (bp) | 1,342.95 |

| | |
|---|---|
| the average exon_number of per gene | 6.97 |
| the average of exon_length (bp) | 269.31 |
| the average of intron_length (bp) | 1,321.07 |
| the total number of exon | 74,587 |
| the total number of intron | 63,891 |
| the total intron length (bp) | 84,404,644 |

**Table S9 Functional annotation of protein coding genes in *K. lacca* genome.**

| Database | Annotated Number | Annotated Percent (%) |
|---|---|---|
| Uniprot | 8,839 | 82.64% |
| Pfam | 8,264 | 77.26% |
| GO | 7,452 | 69.67% |
| KEGG | 5,888 | 55.05% |
| Pathway | 3,373 | 31.54% |
| Interproscan | 9,199 | 86.00% |
| NR | 8,850 | 82.74% |
| Annotation | 9,599 | 89.74% |
| Total | 10,696 | 100% |

**Table S10 The statistical results of non-coding RNA**

| Type | | Copy | Average length (bp) | Total length (bp) | % of genome |
|---|---|---|---|---|---|
| miRNA | | 21 | 79 | 1,663 | 0.000648 |
| tRNA | | 77 | 76 | 5,825 | 0.002270 |
| rRNA | **rRNA** | 57 | 251 | 14,322 | 0.005581 |
| | 18S | 21 | 459 | 9,634 | 0.003754 |
| | 28S | 7 | 144 | 1,009 | 0.000393 |
| | 5.8S | 7 | 158 | 1,106 | 0.000431 |
| | 5S | 22 | 117 | 2,573 | 0.001003 |
| snRNA | **snRNA** | 91 | 128 | 11,604 | 0.004522 |
| | CD-box | 18 | 122 | 2,196 | 0.000856 |
| | HACA-box | 2 | 201 | 402 | 0.000157 |
| | splicing | 70 | 127 | 8,876 | 0.003459 |
| | scaRNA | 1 | 130 | 130 | 0.000051 |

**Table S11 Summary of the predicted protein-coding genes in *K. lacca* genome.**

| Gene set | Number | Average gene length (bp) | Average CDS length (bp) | Average exons per gene | Average intron length (bp) | Average exon length (bp) |
|---|---|---|---|---|---|---|

| | | | | | | | |
|---|---|---|---|---|---|---|---|
| Ab initio | Genscan | 6,500.0 | 23,436.07 | 1,469.3 | 6.42 | 228.79 | 4,051.3 |
| | AUGUSTUS | 14,747.0 | 5,270.22 | 1,338.67 | 6.87 | 194.98 | 670.27 |
| Homology-based | *Diaphorina citri* | 15,935.0 | 7,098.93 | 636.86 | 3.33 | 191.16 | 2,771.6 |
| | *Diuraphis noxia* | 12,179.0 | 8,561.26 | 808.43 | 4.14 | 195.50 | 2,472.81 |
| | *Aphis gossypii* | 11,769.0 | 9,483.99 | 858.1 | 4.40 | 195.16 | 2,539.35 |
| | *Bemisia tabaci* | 11,148.0 | 9,400.94 | 964.23 | 5.00 | 192.94 | 2,110.5 |
| | *Acyrthosiphon pisum* | 13,215.0 | 9,223.38 | 847.4 | 4.22 | 200.83 | 2,601.74 |
| RNA-seq | | 5,210.0 | 13,993.78 | 942.98 | 5.18 | 426.81 | 2,820.63 |
| Integration | | 9,815.0 | 9,746.67 | 1,574.71 | 8.23 | 223.01 | 1,094.88 |
| Final set | | 10,696.0 | 9,769.37 | 1,342.95 | 6.97 | 269.31 | 1,321.07 |

**Table S12** Genes used for gene family clustering in each species.

| Species name | Genes number | Genes in families | Family number | Unique families | Average genes per family |
|---|---|---|---|---|---|
| *Anopheles gambiae* | 9,646 | 9,646 | 7,243 | 997 | 1.33 |
| *Anoplophora glabripennis* | 13,918 | 13,918 | 9,238 | 814 | 1.51 |
| *Acyrthosiphon pisum* | 17,401 | 17,401 | 9,923 | 1,252 | 1.75 |
| *Bombyx mori* | 13,704 | 13,704 | 9,896 | 738 | 1.38 |
| *Bemisia tabaci* | 12,776 | 12,776 | 9,170 | 1,426 | 1.39 |
| *Cimex lectularius* | 11,790 | 11,790 | 9,088 | 1,168 | 1.30 |
| *Drosophila melanogaster* | 13,838 | 13,838 | 10,050 | 2,496 | 1.38 |
| *Diuraphis noxia* | 11,082 | 11,082 | 8,326 | 209 | 1.33 |
| *Danaus plexippus* | 12,684 | 12,684 | 9,555 | 588 | 1.33 |
| *Daphnia pulex* | 27,196 | 27,196 | 13,155 | 6,591 | 2.07 |
| *Ericerus pela* | 13,255 | 13,255 | 9,378 | 2,135 | 1.41 |
| *Kerria lacca* | 10,696 | 10,696 | 7,727 | 702 | 1.25 |
| *Nilaparvata lugens* | 17,347 | 17,347 | 10,511 | 2,195 | 1.65 |
| *Nasonia vitripennis* | 13,539 | 13,539 | 9,162 | 1,492 | 1.48 |
| *Phenacoccus solenopsis* | 7,662 | 7,662 | 5,341 | 441 | 1.43 |
| *Tribolium castaneum* | 12,681 | 12,681 | 9,276 | 798 | 1.37 |

**Table S13** **GC content of HGT adjacent genes**

| Adjacent genes upstream of HGT\|GC content | HGT\|GC content | Adjacent genes downstream of HGT\|GC content |
|---|---|---|
| *Kerria0046200.1*\|0.254 | *Kerria 0046210.1*\|0.278 | *Kerria 0046220.1*\|0.268 |

**Table S14** **FPKM of 11 putative genes of different developmental stages related to Fatty acid synthesis and Terpenoid biosynthesis**

| | Gene ID | L | A1 | A2 | A3 | Functional categories | Pathway |
|---|---|---|---|---|---|---|---|
| TBS1 | gene 0001970.1 | 4.71 | 14.04 | 18.77 | 37.15 | decaprenyl-diphosphate synthase | Terpenoid biosynthesis |
| TBS2 | gene 0043050.1 | 64.10 | 210.03 | 267.67 | 84.26 | | |
| TBS3 | gene 0000520.1 | 9.07 | 14.59 | 14.58 | 51.98 | Farnesyl diohosphate synthase | |
| FAS1 | gene 0044650.1 | 39.33 | 51.28 | 55.38 | 149.41 | acyl-CoA dehydrogenase | Fatty acid synthesis |
| FAS2 | gene 0028390.1 | 18.33 | 28.60 | 30.56 | 111.82 | acyl-CoA synthetase | |
| FAS3 | gene 0074180.1 | 32.65 | 68.47 | 37.75 | 33.36 | aldehyde dehydrogenase | |
| FAS4 | gene 0012900.1 | 3.36 | 10.82 | 150.38 | 5.98 | fatty acid elongase | |
| FAS5 | gene 0019660.1 | 6.61 | 69.82 | 82.60 | 11.63 | | |
| FAS6 | gene 0019670.1 | 17.72 | 26.38 | 227.88 | 379.24 | | |
| FAS7 | gene 0008080.1 | 99.70 | 126.99 | 110.13 | 244.70 | S-malonyltransferase | |
| FAS8 | gene 0071240.1 | 8.02 | 12.82 | 22.59 | 33.12 | very long-chain-fatty-acid--CoA ligase | |

**Table S15** **FPKM of 14 putative genes of different tissues related to Fatty acid synthesis and Terpenoid biosynthesis**

| | ID | AT | BH | BW | MG | Functional categories | Pathway |
|---|---|---|---|---|---|---|---|
| TBS-1 | gene 0038380.1 | 41.88 | 48.08 | 96.65 | 6.62 | dehydrodolichyl diphosphate synthase | Terpenoid biosynthesis |
| TBS-2 | gene 0013810.1 | 53.56 | 118.05 | 275.74 | 47.34 | | |

| | | | | | | | |
|---|---|---|---|---|---|---|---|
| TBS-3 | gene 0005780.1 | 25.21 | 58.40 | 231.94 | 8.04 | diphosphomevalonate decarboxylase | |
| TBS-4 | gene 0000530.1 | 34.57 | 87.36 | 299.07 | 10.52 | | |
| TBS-5 | gene 0019400.1 | 173.14 | 440.11 | 1780.33 | 28.76 | farnesyl diphosphate synthase | |
| TBS-6 | gene 0038900.1 | 4.40 | 6.62 | 47.60 | 2.62 | | |
| TBS-7 | gene 0046210.1 | 171.01 | 315.47 | 1168.25 | 20.09 | isoprenyl transferase | |
| FAS-1 | novel741 | 41.44 | 72.58 | 87.10 | 6.35 | 3-hydroxyacyl-CoA dehydratase | |
| FAS-2 | gene 0018240.1 | 393.82 | 756.68 | 564.45 | 83.19 | | |
| FAS-3 | gene 0087400.1 | 180.52 | 232.78 | 261.75 | 131.42 | Acetyl-CoA C-myristoyltransferase | |
| FAS-4 | gene 0019590.1 | 1651.29 | 2396.34 | 6918.93 | 959.97 | fatty acid elongase | Fatty acid synthesis |
| FAS-5 | gene 0029260.1 | 6.66 | 14.68 | 80.20 | 1.55 | | |
| FAS-6 | gene 0038400.1 | 243.98 | 208.39 | 432.53 | 1.57 | long-chain-fatty-acid--CoA ligase | |
| FAS-7 | gene 0042820.1 | 314.15 | 181.59 | 572.79 | 11.73 | | |

**Table S16 The quantitative PCR primers of 25 candidate genes in *K. lacca***

| Gene Name | primer sequence | Tm | Gene length |
|---|---|---|---|
| TBS1 | F:GAATTGCCTGAGTGAATGTA | 49 | 112 |
| | R:CGATGATGGTGAATCTGTAG | 49.1 | |
| TBS2 | F:GCTCTTCGTGTAATAGTCTTG | 50.2 | 139 |
| | R:TCATCACTCAATAGCCATCT | 49.1 | |
| TBS3 | F:TGGAACTGCTGTGGCTAA | 51 | 149 |
| | R:CACCTGGTACATTGTATTGAAG | 51 | |
| FAS1 | F:AGAATTGGTATCGCATCACA | 50.5 | 151 |
| | R:CGAGACGAAGAGCCATATC | 50.6 | |
| FAS2 | F:GGACAGGAACGATTGAAGA | 49.9 | 121 |
| | R:CGCCAAGTGTAGCATAGAT | 50.2 | |
| FAS3 | F:CAGAAGGAATCGGAGAAGT | 49.3 | 183 |
| | R:CATAAACGGCAACAGGAAA | 49.4 | |
| FAS4 | F:GCACCTATACCATCATTCG | 48.5 | 124 |
| | R:ATAGCAGTACATCACCACAT | 49.4 | |
| FAS5 | F:ATGCTTACTCCGCTATATGG | 50.2 | 103 |
| | R:AACTGGTCGTATTCGCTTAT | 50.3 | |
| FAS6 | F:TTATAGTCGGCGGTATAGGA | 50.3 | 168 |
| | R:GATGAACTCTGTGATCTCTTAC | 50 | |

| | | | |
|---|---|---|---|
| FAS7 | F:GGAGAATTAACCGCACTGA | 50.3 | 119 |
| | R:GCCATACCACCATTAGACAT | 50.5 | |
| FAS8 | F:CAGAACAGGCAGCGTATT | 50.2 | 194 |
| | R:GCGAACATCTCTTGTAGCA | 50.8 | |
| TBS-1 | F:GGATACTACACAGTCGTCTT | 49.7 | 132 |
| | R:CGATTACCGTCCATTACAAT | 49 | |
| TBS-2 | F:CAGAAGCAGCATTAGATGAC | 50 | 111 |
| | R:GCCAAGGAGATAATCCATAAG | 49.5 | |
| TBS-3 | F:TTCCATAAGTGCGACTCTAT | 49.2 | 105 |
| | R:TCCTCTATTCCATTCAACCA | 48.9 | |
| TBS-4 | F:TACCGACAGCAATAGCAAT | 49.6 | 198 |
| | R:CAGCCACTATAAGCCATGT | 50 | |
| TBS-5 | F:TATTCTGTGTTAGCGGATGA | 49.6 | 100 |
| | R:CATAAGATGCTACTGGACTTC | 49.5 | |
| TBS-6 | F:TGACGAATGTTACGGTTGTA | 50.5 | 156 |
| | R:TGCGGAAGACCTCTTGATA | 50.8 | |
| TBS-7 | F:GGAGCGGTTATTCAATTAGC | 50.3 | 141 |
| | R:AACTCCGTGGACATTCATC | 50.3 | |
| FAS-1 | F:GTGTCGGTGGAGAATTATTAG | 49.8 | 179 |
| | R:TCTGCGTTGAGTGAACAT | 49.3 | |
| FAS-2 | F:GAATTGCCGAACCAACTAA | 49 | 145 |
| | R:CTGTAGTGATCTGAGTTCCA | 49.5 | |
| FAS-3 | F:CTACAGGTTCTTCAGCATTAC | 50 | 164 |
| | R:TTCAGCCATTACTTCCACAT | 50.2 | |
| FAS-4 | F:CATTCCAGAATCACGCATT | 49.2 | 130 |
| | R:CGACACAATAACCATCCTAC | 49.3 | |
| FAS-5 | F:CGATGACGAAGGATGGTTA | 49.9 | 179 |
| | R:TTAGAACTGCTTGACTGACA | 49.9 | |
| FAS-6 | F:GACTGACGAAGAAGGATGG | 50.5 | 175 |
| | R:GGCTTGACTGAGAATTGGT | 50.5 | |
| FAS-7 | F:ATTCAGCGTTAGCGACAA | 49.8 | 194 |
| | R:GATTCCTCCTGCCATTATTG | 49.5 | |
| 28S rRNA | F:TACTGACTGCCCGGATTC | 50 | 134 |
| | R:AATACGACCACCGAGACC | 50 | |
| RPL13 | F:CATCGTCGTCGTAATAAGTC | 50 | 135 |
| | R:TTCTTCTTCCGTAGCTTCTC | 50 | |
